# Systemic autoimmune disease patients’ blood immunome reveals specificities and commonalities among different diagnostic entities

**DOI:** 10.1101/2024.05.27.594621

**Authors:** Paulina Rybakowska, Sofie Van Gassen, Guillermo Barturen, Carlos Pérez Sánchez, Alejandro Ibáñez-Costa, Nieves Varela, Rafaela Ortega Castro, Concepción Fernández-Roldán, Inmaculada Jiménez-Moleón, Norberto Ortego, Enrique Raya, Rocío Aguilar Quesada, Chary López-Pedrera, Eduardo Collantes, Yvan Saeys, Concepción Marañón, Marta E. Alarcón-Riquelme

**Affiliations:** Pfizer-University of Granada-Junta de Andalucía Centre for Genomics and Oncological Research (GENYO), PTS, Granada, Spain; Department of Applied Mathematics, Computer Science and Statistics, Ghent University, Gent, Belgium; Data Mining and Modeling for Biomedicine, VIB Center for Inflammation Research, Gent, Belgium; Maimonides Institute for Research in Biomedicine of Cordoba (IMIBIC)/Reina Sofia University Hospital/University of Córdoba, Spain; Servicio de Medicina Interna. Unidad de Enfermedades Autoinmunes Sistémicas. Departamento de Medicina, Universidad de Granada, Spain. Hospital Universitario San Cecilio, PTS. Granada, Spain; Servicio de Reumatología. Hospital Universitario San Cecilio, PTS, Granada, Spain; Biobanco del Sistema Sanitario Público de Andalucía, Andalusian Public Health System Biobank, Granada, Spain; Institute for Environmental Medicine, Karolinska Institutet, Stockholm, Sweden

**Author notes:** These authors contributed equally (Concepción Marañón, Marta E. Alarcón-Riquelme).

**Keywords:** Systemic autoimmune diseases, autoimmunity, mass cytometry, cytometry by time-of-flight, immunophenotyping, patients’ stratification, biomarkers

## Abstract

1

**Background:** Systemic autoimmune diseases (SADs) are characterized by internal heterogeneity, overlapping clinical symptoms, and shared molecular pathways. Therefore, they are difficult to diagnose and new tools allowing precise diagnosis are needed. Molecular-based reclassification studies enable to find patterns in a diagnosis-independent way.

**Objective:** To evaluate the possibility of using high-content immunophenotyping for detecting patient subgroups in the context of precise treatment.

**Methods:** Whole blood high-content immunophenotyping of 101 patients with 7 systemic autoimmune diseases and 22 controls was performed using 36-plex mass cytometry panel. Patients were compared across diagnostic entities and re-classified using Monte Carlo reference-based consensus clustering. Levels of 45-plex multiplexed cytokine were measured and used for cluster characterization.

**Results:** Differential analysis by diagnosis did not reveal any disease-specific pattern in the cellular compositions and phenotypes but rather their relative similarities. Accordingly, patients were classified into phenotypically distinct groups composed of different diagnostic entities sharing common immunophenotypes and cytokine signatures. These features were mainly based on granulocyte activation and CD38 expression in discrete lymphocyte populations and were related to Th17 or IFN-dependent cytokines.

**Conclusions:** Our data indicate that specific individuals could potentially benefit from the same line of treatment independently of their diagnosis and emphasize the possibility of using immunophenotyping as a stratification tool in precision rheumatology.

**Graphical abstract:** 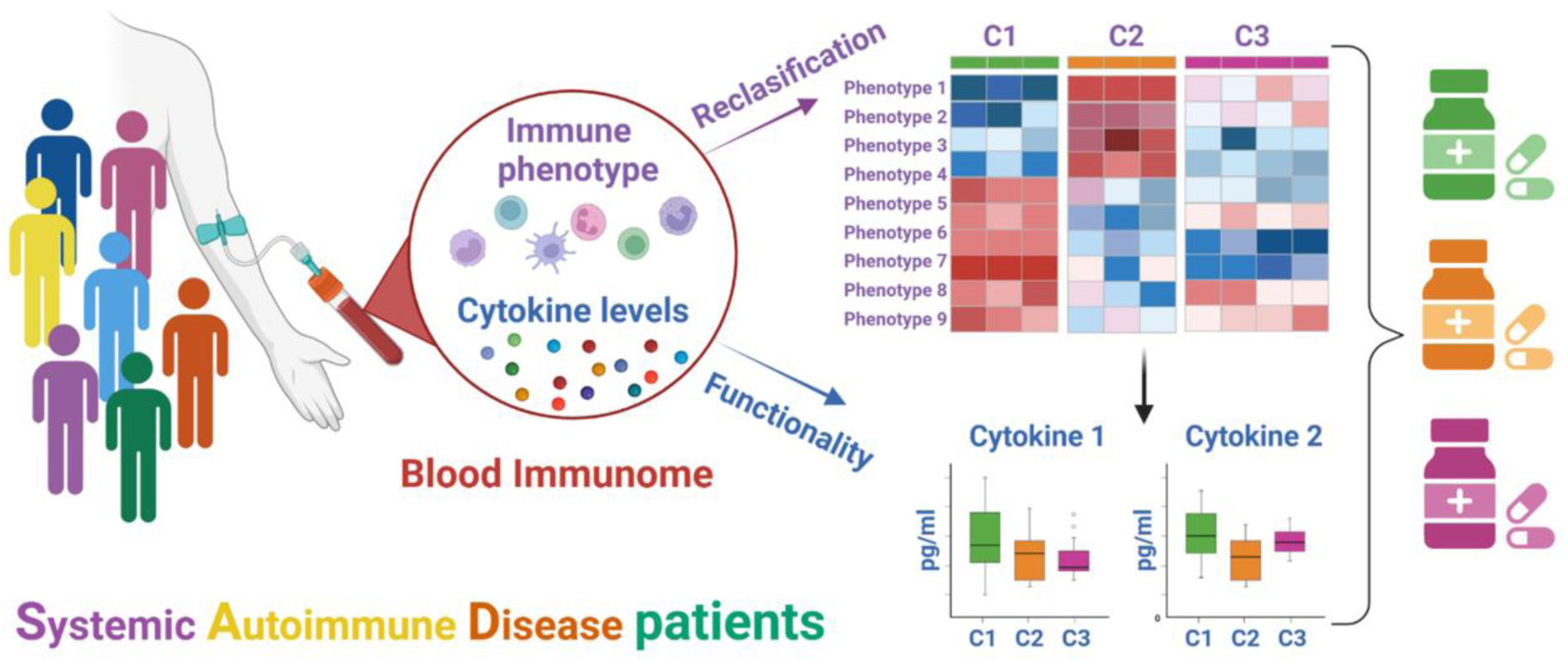

**Key messages:**

- Whole blood immmunophenotyping could be used to stratify systemic autoimmune patients, thus it is a useful tool in precision medicine.
- Patients’ groups could benefit from the same line of treatment.

## 5 Introduction

Systemic autoimmune diseases (SADs) are diagnosed using different clinical and laboratory criteria^1^. Due to the high internal heterogeneity within each clinical entity and overlapping symptoms between diseases, differential diagnosis of SADs is complex. Accordingly, the time from disease onset to diagnosis can take years^2^, leading to a poorer prognosis. A fraction of the patients can be classified as mixed connective tissue disease (MCTD) which is still a controversial disease entity^3,4^ or even as undifferentiated connective tissue disease (UCTD), a group that comprises those patients without a clear clinical picture, and hence without a diagnosis and appropriate treatment^2^.

Such heterogeneity and the overlaps between diseases can be observed at molecular, genetic and cellular levels. For instance, the presence of anti-SSA and SSB antibodies and their genetic association with HLA class II allele *DRB1*0301* was found in both systemic lupus erythematosus (SLE) and Sjögren’s Syndrome (SJS) patients^5,6^. The cell frequency of circulating leukocytes can be decreased or increased in different cohorts of the same disease, which can be due to different experimental settings, but also to different endotypes. When comparing immune cell composition between diseases, not many differentially expressed features could be found^7,8^, suggesting that immune cell landscapes between patients with different SADs could be shared in some extent. Additionally, the current treatments although effective, show high variability in the responses observed, and groups of non-responders are still elevated, indicating an inadequate molecular fit between the treatments and the actual pathogenesis.

In response to these problems the large-scale study PRECISESADS included 7 different SADs (SLE, SJS, UCTD, MCTD, Systemic Sclerosis (SSC), Rheumatoid Arthritis (RA), and primary anti-phospholipid syndrome (PAPS)) and showed patient classification into 4 different molecular groups based on transcriptomic and epigenomic data^9^. The enrichment of circulating cell populations in different clusters was also shown e.g., neutrophil enrichment in the inflammatory clusters. However, no further cellular characterization was performed due to the low resolution in characterization of circulating populations. Although these results confirm commonalities behind different SADs, due to application of high-dimensional OMICS technologies, they are difficult, expensive, and time-consuming to introduce into clinical practice. Thus, alternative techniques already used in the clinical hospitals, like cytometry based immunophenotyping could be of interest.

Until recently, 2 (SJS, SLE)^8^ and 3 diseases (SLE, SJS, systemic sclerosis (SSC))^7^ had been compared using high-throughput immunophenotyping. As expected, both studies showed similar molecular mechanisms between different groups of patients. Additionally, in both experimental groups only peripheral blood mononuclear cells (PBMCs) were considered, and no functional markers were measured. In another study using flow cytometry, the diseases SLE, SJS, RA, and SSC were compared, and a common inflammatory phenotype was found across them, but only B and T cells antibody panels were used, limiting the scope of the findings^10^.

Keeping in mind the above-mentioned limitations and the utility of immunophenotyping, we aimed at applying high content cytometry to perform whole blood deep-phenotyping, including measures of cell frequency and functional marker expression levels. The goal was to compare patients from multiple SADs using a sub study from PRECISESADS cross-sectional cohort, and additionally reclassify them based on their immunological landscape. A deeper look into immune composition and immune responses of different SADs can give a better understanding of ongoing immunopathology in each patient and get us closer to precision medicine in rheumatic diseases.

## 6 Materials and Methods

### 6.1 Study participants

Eligible patients were aged 18 or older and diagnosed as having one of the following SADs: SLE (n = 23), SJS (n = 22), SSC (n = 18), RA (n = 15), MCTD (n = 6), PAPS (n = 4). Patients without any diagnosis but with manifestations clearly associated with the clinical diagnoses were also recruited and termed UCTD (n = 13). Eligible healthy controls (CTRs, n = 22) were matched on the projected and expected profile of patients in terms of age. Individuals were recruited in four recruitment centers (Department of Rheumatology and Systemic diseases, Clinical Hospital San Cecilio, Granada, Spain; Andalusian Health System Biobank, Granada, Spain; Department of Rheumatology, Hospital Reina Sofia, Córdoba, Spain). Each patient was diagnosed according to the prevailing international classification or diagnostic criteria established for each of the SADs^11–16^. For UCTD we considered patients with clinical features of SADs not fulfilling any of the SADs criteria (RA, SSc, pSjS, SLE, PAPs, MCTD), or any other SADs criteria for at least 2 years, with presence of antinuclear antibodies (ANA) ≥ 1:160 with or without other specific autoantibodies; patients fulfilling 3 out of 11 SLE ACR classification criteria and patients with early systemic sclerosis^17^ were not classified as UCTD. All donors signed an informed consent according to the ethical protocol of the Andalusian Biobank and the PRECISESADS project (ClinicalTrial identifier NCT02890121). The project protocol was approved by the Ethical Committee of Centro Granada (CEI-Granada), Spain and Hospital Universitario Reina Sofía Córdoba, Spain, according to the Helsinki declaration of 1975, as revised in 2013. The main exclusion criteria can be found in previously published work^9^. Demographic information and drug prescriptions are given in Table S1.

### 6.2 Sample processing for deep-phenotyping study using mass cytometry

Samples were processed in two research centers in Granada (GENYO, GRA) and Córdoba (IMIBIC, COR), following the SOP prepared for the PRECISESADS project as described before^18^. Briefly, 500 µl of blood collected in EDTA-K3 vacutainer tubes was stained with live/dead reagent (CisPt, 5μM) for 10 min, RT. Blood cells were fixed for 10 min with proteomic stabilizer (PROT, SmartTube) and frozen at -80°C until staining. A reference sample was prepared to track and correct batch effect in this multibatch study as described before^19^. The reference sample consisted of 6 ml of whole blood from a single donor, was fixed with 7.2 ml of PROT, as for donor samples. Aliquoted and stored at -80°C until the time of staining.

### 6.3 Sample staining and acquisition on CyTOF/Helios

Samples were assigned to 9 experimental batches with even distribution of clinical groups. Each batch consisted of 15 samples + 1 reference sample and was stained with an aliquots of a frozen antibody cocktail (Table S2), prepared as described^20^. Most of the antibodies were obtained in a labeled form from Standard BioTools. Alternatively, purified antibodies were conjugated using MaxPar Metal-labeling kits (Standard BioTools), following the vendor protocol.

Blood samples were thawed, lysed and stained for surface markers as described^21^. Briefly, 1.5*10^6^ cells/sample were used for the barcoding with Cell-ID 20-Plex Pd Barcoding Kit (Standard BioTools). After barcoding, samples were washed with cell staining buffer (CSB, Standard Biotools), pooled and surface antigens were stained with the antibody cocktail in CSB, for 30 min at 4°C. After washing with CSB, cells were stained with Iridium (Ir, 5 μM, Standard Biotools) for 20 min, at RT in Fix and Perm Buffer (Standard BioTools), washed with CSB and left O/N in 2% formaldehyde (PFA) (Thermo Fisher Scientific) at 4°C.

The following day the acquisition was performed in aliquots to avoid prolonged cell exposure to water as described^21^. Samples were acquired with EQ Four Element Calibration Beads (Standard BioTools) in a CyTOF2/Helios device using a narrow bore sample injector. The flow rate was set below 400 events/s and each aliquot was acquired for no longer than 2h.

### 6.4 CyTOF data preprocessing and cell phenotype extraction

Quality control, data preprocessing and normalization using calibration beads and the reference sample were performed as described before^19^. Granulocytes (GRAN) and PBMC were analyzed separately. PBMC and granulocyte gating was done using the ratio computed for CD15 and CD45 markers, see representative gating in Figure S1A. For the PBMC compartment FlowSOM was built^22^ using PBMC-clustering markers (Table S2). An aggregated file was created to construct the FlowSOM tree by randomly subsetting 25,000 PBMC per fcs file, generating a file with 3.075*106 cells in total. A grid of 20x20 clusters was used, and 45 metaclusters (P1-P45) were identified using consensus clustering. Each individual file was then mapped to the aggregated FlowSOM and cell frequencies and median signal intensities (MSI) for functional markers (see Table S2) were calculated for each P. To check the quality of the clustering, the aggregated file was manually gated using FlowJo 10.0.7 (Figure S1), and the F1-measure together with weighted purity scores was calculated as described (1). The manual labels Figure S1 for each P were assigned using the cell population representing the majority of the cells (enriched cell populations) included in a particular P (Figure 1A, B). Enriched subset names are used across the manuscript in parenthesis, together with the metacluster numbers. The metacluster defined as Unlabeled (grey) were those not assigned to any population during manual gating (Figure 1). To identify granulocyte cell subsets, PhenoGraph^23^ was used together with granulocytes-clustering markers (Table S2). To run Phenograph the data were first aggregated taking 2,500 granulocytes per fcs file, building an fcs file with 307,500 granulocytes in total. Next, clustering was run on the aggregated file with the parameter k (defining the number of the nearest neighbors) set to 100 as described before^24^, in total 17 clusters were detected. Similarly to FlowSOM setting, the ConsensusClusterPlus^25^ was used to perform consensus clustering and finally 8 metaclusters (G1-G8) were identified (Figure 1C, D). Cell frequencies and MSI for functional markers (Table S2) were calculated for each G using the aggregated file.

**Figure 1.**
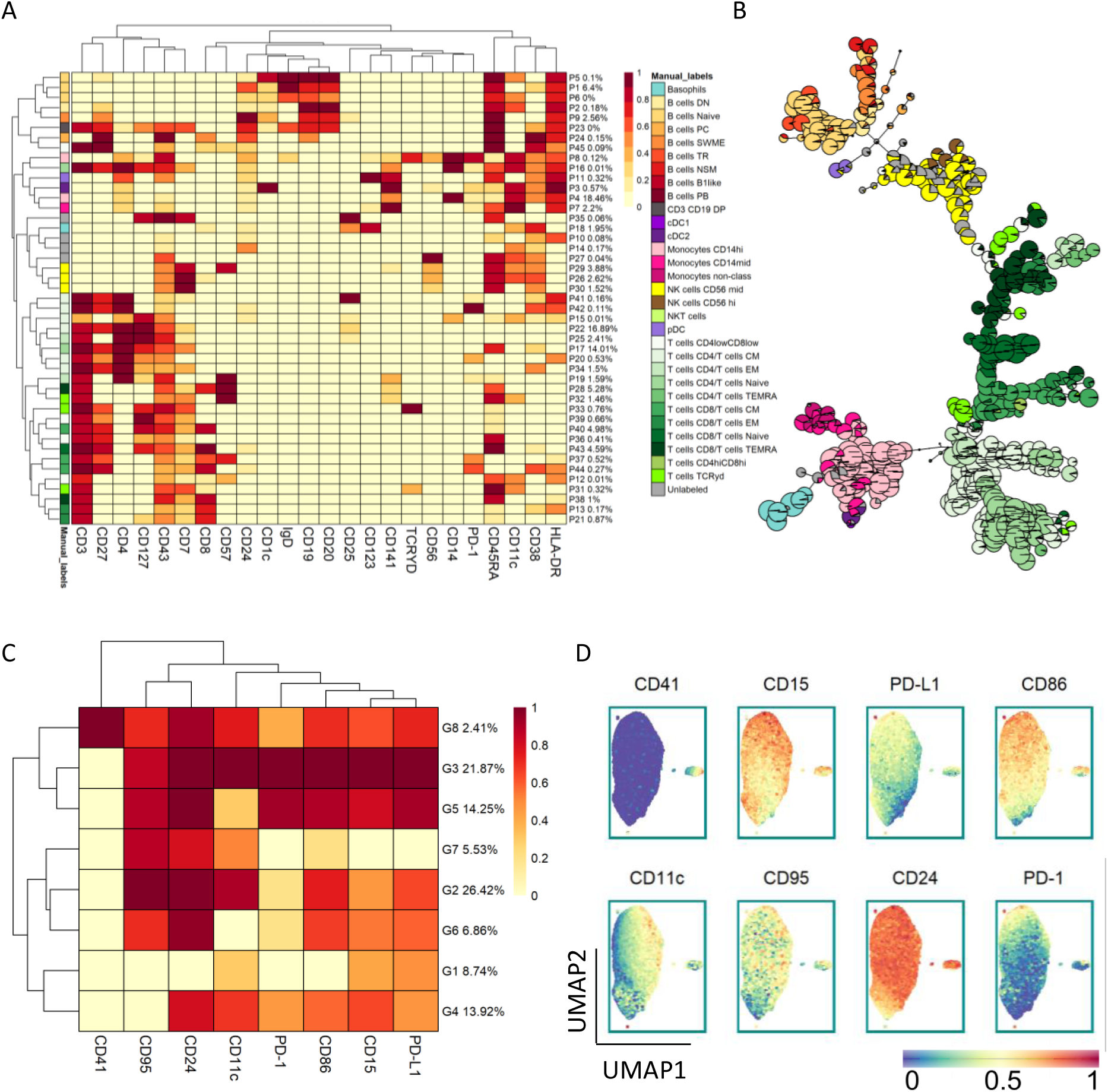
FlowSOM (PBMC) and Phenograph (GRAN) clustering. (A) Heatmap represents median intensities for clustering markers across the 45 FlowSOM metaclusters. The color intensity in the heatmap represents the median of the arcsinh, 0-1 transformed marker expression of the cells included in the metacluster in the aggregated file. The colors assigned to the metaclusters correspond to the manual gating labels and are the same throughout the figures corresponding to PBMC analysis. The relative size of each metacluster is indicated in the heatmaps’ row names. The dendrograms represent the hierarchical clustering using Euclidian distance and average linkage. (B) Manual labels overlaid on FlowSOM tree. Pie charts indicate the percentage of the cells represented by the node falling in the manual gates. A different color is assigned to each label and the manual annotation can be found in the legend. (C) Heatmap for granulocytes represented as in B. (D) Granulocytes’ clustering visualization by UMAP. UMAP was built using the aggregated file and clustering markers used in Phenograph analysis. Cells are colored according to the expression level of clustering markers. The expression is 0-1 transformed.

The P and G metaclusters with a median cell count lower than 50 cells were removed from further analysis. Similarly, features consisting in markers with low, median expression (arcsine transformed MSI) < 1 in specific metacluster were also removed. CD28 marker was excluded from the analysis, since it generated a strong batch effect despite file normalization. In total we extracted 159 features. Total number of granulocytes or PBMC was used as the reference population to quantify G and P frequencies, respectively. The probes used for the clustering and functional markers can be found in Table S2.

Cell clustering and analysis were performed using R-implemented algorithms. Data were visualized using aggregated files. Additionally a UMAP visualization^26^ was performed using *uwot* package and a random subset of 2,500 cells per file. Metacluster frequencies or MSI were represented on two-dimensional maps using *ggplot2* package. Heatmaps were generated using *pheatmap*.

### 6.5 Sample processing and quantification for cytokine/chemokine study in plasma

Two hundred fifty μl of blood obtained in EDTA-K3 tube was diluted 1:1 with RPMI (Gibco) and incubated at 37°C, 5-7% CO_2_ for 24h. Next, samples were spun at 800 g for 5 min 4°C, and the supernatants (diluted plasma) were collected and stored frozen at -80°C, until the analysis by Luminex. Forty-five analytes were measured using diluted plasma and Human XL Cytokine Discovery Premixed Kit (R&D System). Samples were thawed on ice and diluted 1:2 with calibration diluent. The assay was performed adapting the protocol recommended by the manufacturer, by using half volume of the samples and reagents, and incubating the beads with plasma O/N at 4°C. At least 50 beads per analyte were acquired using a Luminex 200 device. The concentration of each protein was calculated using the corresponding standard curve and the Bio-Plex Manager v6.0 software. The standard curves were created using a 4 or 5-parametere logistic curve fit and were expressed as pg/ml.

The beads with less than 25 counts were removed from the analysis of this sample. Cytokines for which no expression was detected in 90% of the samples (IL-17A, IL-3, IL-7, IL-4, IL-5) were removed from the analysis. The imputation for out-of-range values (OOR) was performed as follows: for high detection range the value was imputed as the maximum value plus 50%; for data in the low detection range, we imputed half of the minimum value min.

### 6.6 Quantification and statistical analysis

Differential analysis for all the diagnostic groups was performed using Kruskal-Wallis (K-W) and Dunn *post-hoc* test. The false discovery date (FDR) method was used to correct for multiple testing. Agglomerative hierarchical clustering was performed using ward.D2 linkage.

To select features for patient reclassification, a linear model was performed comparing CTRs and SADs using extracted features that passed quality control. The features with p-values ≤ 0.01 were selected for Monte Carlo reference-based consensus clustering (M3C) using the partition around medoids (PAM) method^27^ on SADs samples. One thousand iterations and 90% of the data were re-sampled with each iteration. The optimal number of clusters was selected using the lowest p value along the range of clusters (1-20).

To verify the association of the features with medication, a Mann Whitney (M-W) test was performed between treated (Tx), non-treated (NTx) and healthy controls (CTR), and correction for multiple testing (FDR) was applied. The differences in cytokine production between clusters were calculated by K-W test, followed by Dunn test, as described above. The differences in clinical parameters across clusters were calculated using Fisher exact test (F-T), and K-W was used for age and disease duration. K-W and M-W analysis were performed using *rstatix* package. Plotting was done using *pheatmap* and *ggplot2* packages. All analyses were performed in R, except cytokine visualization across clusters, which was done using GraphPad Prism v6.01.

## 7 Results

### 7.1 The phenotype of circulating cells reveals similarities between diagnostic groups

To deeply characterized circulating immune cell populations we performed automated, unsupervised clustering. We obtained 45 and 7 distinct metaclusters in the PBMC and GRAN compartments, using FlowSOM and Phenograph, respectively (Figure 1 and Table S2 for markers used). In total, we analyzed 159 features including cell frequencies and MSI. PBMC metaclusters were labeled using the expression of clustering markers (Figure 1A). When comparing with classical manual gating (Figure S1) we found a good correspondence with an average purity per cluster of 0.91 (data not shown). When grouping the clusters into metaclusters, the average purity was 0.84 (data not shown) and there was a clear match with the manual populations (e.g., P3 corresponds with CD1c^+^ cDC, Figure 1A). This translates into an F1-measure equal to 0.81 (data not shown), confirming good metacluster results obtained by FlowSOM. Because of this, we use hereon manual labels as metacluster descriptors, keeping in mind that they are enriched in, rather than composed of, pure cell types. Using the unsupervised clustering algorithm Phenograph we detected 7 neutrophil subpopulations: for instance, CD11c^lo^ CD24^lo^ known as freshly released from bone marrow and CD11c^hi^ and CD24^hi^ called aged neutrophils^28,29^. Due to the less defined phenotype of the granulocyte compartment we did not obtain manual labels for these cells. Curiously, we detected a G8 population that was positive for canonical platelets’ marker CD41^+^, suggesting the platelets-associated with neutrophils^30^. Because of the difficulty in preserving eosinophils-specific markers upon fixation, we could not separate eosinophils from neutrophils, thus these cells were analyzed together. Therefore, the interpretation of CD11c^low^ clusters that could potentially contain eosinophils should be carefully taken.

As a first approach, we explored the differences in the frequency and expression of functional markers in relation to the clinical diagnosis. The exploratory analysis using UMAP and density plots showed evident differences in the cell distribution across different diagnoses. For example, an increase in monocyte frequency can be observed in SLE and RA (Figure 2A). Differences in the MSI of several functional markers were also noticed, for example CD38 expression in different NK cell populations and PD-1 in granulocyte subset G3, Figure 2A.

**Figure 2.**
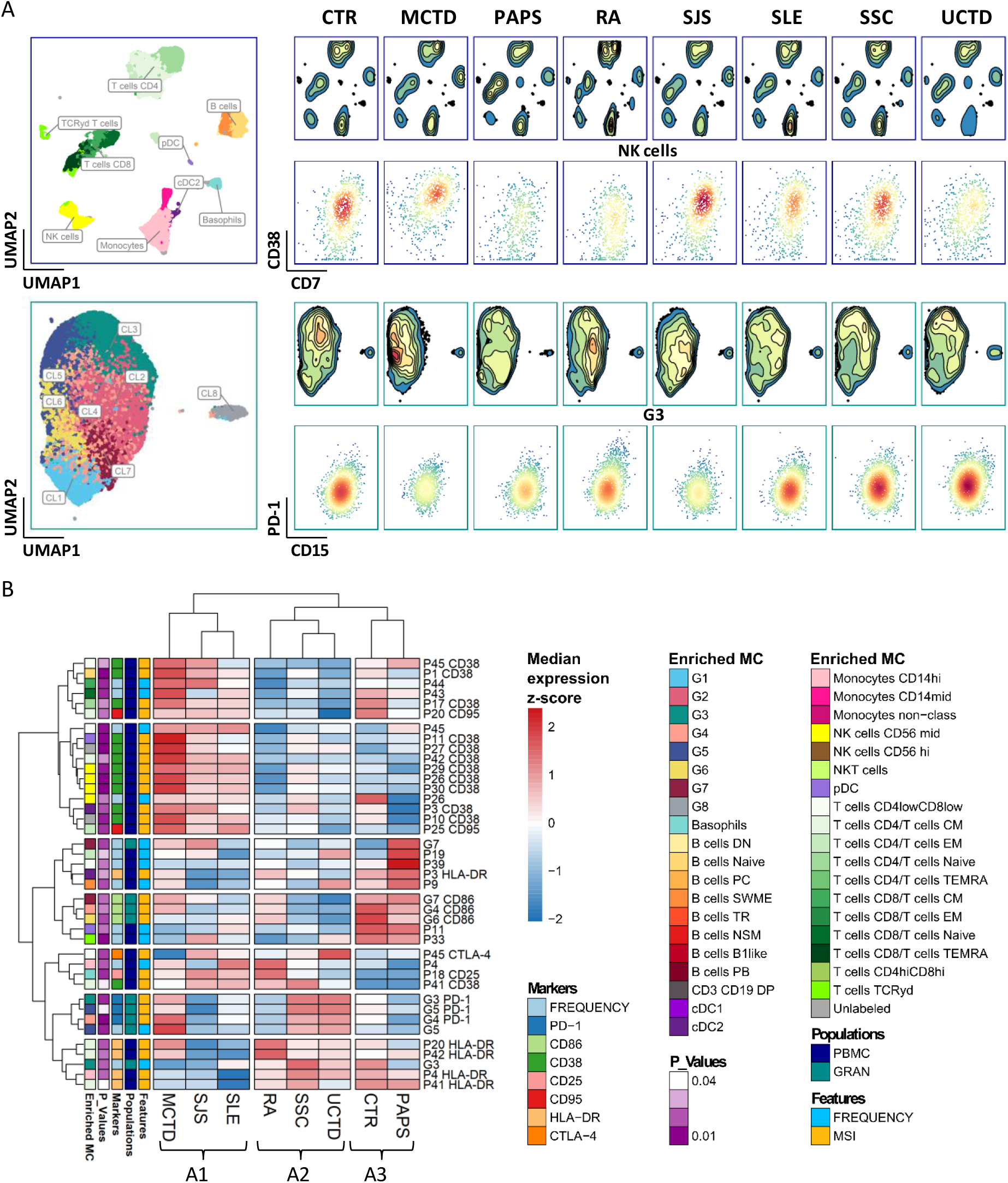
The immunophenotype of circulating cells shows similarities between different disease entities. (A) UMAP was built using the aggregated file and arcsine-transformed expression of clustering markers in the PBMC (green) and GRAN (blue) compartment. Equal numbers of randomly selected cells (10,000 per diagnosis) from each disease are represented. The map is colored by manual or cluster labels (left in PBMC and GRAN, respectively); by the density (top, right in PBCM and GRAN); or biaxial plots are showed for selected subpopulations NK cells for PBMC and G3 cluster for GRAN (bottom, right). (B) Heatmap represents features with significant values in the K-W. The median for each feature was calculated for each disease label. Color scale represents the z-score. Hierarchical clustering with Euclidian distance and ward.2D linkage was performed, and 3 groups of diseases and 7 groups of features were identified. The redder the color, the denser the region or the highest is the expression.

To obtain a deeper visualization into alterations between patient groups we performed a K-W analysis, followed by *post-hoc* Dunn test. In total 40 features (27 for MSI and 13 for cell frequencies) were identified as statistically significant between diagnostic groups. However, no disease specific features were detected (see Figure S2 and Figure S3). Using the per diagnosis median of statistically significant features we performed clustering analysis. Three main groups of diseases (A1, A2, A3) were found (Figure 2B). Group A1 gathered MCTD, SJS and SLE. Within this group, SJS and SLE patients were more related to each other, as already described^8^. Group A2 assembled RA, UCTD and SSC, where more similarities were observed between SSC and UCTD than with RA. CTRs were grouped with PAPS in group A3. The main feature separating groups A1 and A2 from CTR (group A3) were CD38 and HLA-DR expression in specific PBMC populations (P), followed by CD86 and PD-1 expression in granulocyte metaclusters (G). Frequencies of P39, P45 (CD4^low^CD8^low^), P11 (pDC), P4 (CD14^hi^ monocytes), and G3 were also crucial in the formation of the groups.

Group A1 was characterized by higher CD38 expression in the PBMC compartment mainly in P29, P26, P27, P30 (NK cell subsets), P3, P11 (cDC CD1c^+^ and pDC respectively). The strongest CD38 expression was observed in MCTD especially in P1 and P45 (Naïve B cells, CD4^low^CD8^low^ T cells, respectively) and the lowest in patients from group A2. It is worth mentioning that dysregulated CD38 expression on various cell types was already reported in SLE^31^ and a successful treatment using anti-CD38 in refractory SLE was reported^32^. Additionally, lower HLA-DR expression in several populations was observed in SJS and SLE patients compared to the other two groups. This low level was noticed in P20, P41, P42 representing subsets of CD4^+^ memory T cells and P3 and P4 representing cDCs, CD1c^+^, and CD14^hi^ monocytes, respectively. P4 and P41 share similar HLA-DR levels across the three diagnoses in A1. High PD-1 expression in multiple granulocyte metaclusters differentiated UCTD and SSC from RA, SJS and SLE patients, however PD-1 grouped them closer to MCTD. The levels of CD95 and CD86 expression were similar for group A2 and A1 across multiple granulocyte populations and differed from that in group A3. Interestingly, CTLA-4 expression was high in P41 (CD4^+^ T regs) in UCTD and SJS patients and the expression of the activation marker CD25 was elevated in P18 (basophils) in multiple diseases (RA, SLE and SJS) suggesting cell activation. Group A1 and A2 had lower levels of circulating P11 (pDC) and P33 (TCRƴδ T cells) compared to A3, which was already described for various SADs^7,33^.

These results show that patients group together based on cellular phenotypes. No disease-specific biomarkers could be selected, underscoring the common mechanisms shared by different SADs.

### 7.2 The phenotype of circulating cells stratifies SADs patients into 3 diagnosis-independent clusters

To find the common mechanisms across SADs we performed differential analysis between SADs and CTRs using a linear model. Fourteen features were significantly different between SADs and CTRs, in both, GRAN and PBMC compartments (Figure S4A). In granulocytes, CD86 and PD-L1 downregulation and CD95 upregulation was observed across various populations. In PBMC, CD38 upregulation was observed in P27 (Unlabeled) and P41 (memory CD4 T cells), and CD25 upregulation in P18 (basophils). Using these features and the M3C method, we classified individuals into 3 robust clusters (C1-C3) (Figure 3 and Figure S4B) and visualized them by heatmap and UMAP (Figure 3A and B, left, respectively).

**Figure 3.**
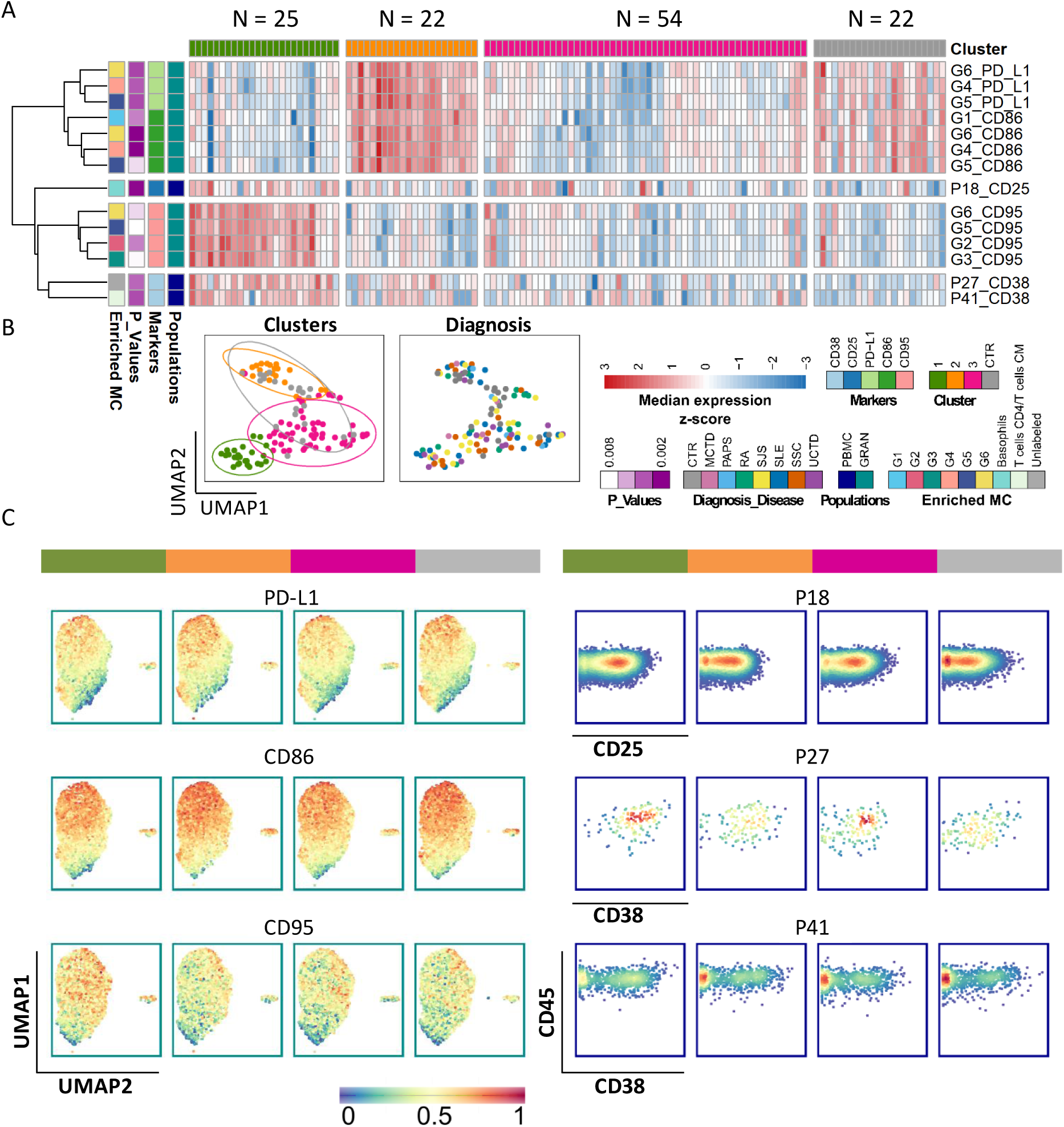
Patients fall into phenotypically distinctive groups based on differentially represented circulating cells. (A) The heatmap represents scaled values (z-scores) for each selected feature across all individuals. In rows, features selected as being differentially expressed between SADs patients (N = 101) and CTRs (control individuals, N = 22) are shown together with the annotation on the left. The annotation colors represent the manual or metacluster label for PBMC and GRAN respectively. In the columns, each individual is plotted together with color-coded cluster assignment. Three clusters were identified (C1 – green, C2 – orange, C3 – pink) by M3C. Additionally, CTRs are shown as a reference in gray. (B) Clustering representation by UMAP analysis colored by cluster assignment, including CTRs (left) or by diagnosis (right). (C) representation of marker expressions by UMAP in GRAN (left) and by biaxial plots in PBMC (right). In PBMC 3 selected metaclusters are presented. CD45 is plotted on the y-axis and subsequent markers on the x-axis.

Clusters, C1 and C2 were opposite to each other, C2 being more similar to CTR. C3 represented the transitional phenotype between C1 and C2, as can be seen in the UMAP visualization, Figure 3B. Cluster C1 was characterized by higher expression of CD95 (Fas receptor) across various granulocyte populations and higher level of the activation markers CD38 and CD25 in specific populations of the PBMC compartment. On the other hand, C2 showed elevated expression of proteins belonging to the B7 family, CD86 and PD-L1 in several granulocyte populations. These proteins are known to be involved in inter-cellular regulation^34,35^.

High level of CD95 in cluster C1 was mostly observed in G2, G3, mature CD11c^+^ granulocytes and G5, G6 immature CD11c^-^ granulocytes. The up regulation of the activation markers CD25 and CD38 in cluster C1 was seen for P18 and P27, P41, respectively (Figure 3 A and C). The phenotypes of these cells were verified using biaxal plots (Figure S5). Showing that P18 cells were basophils, P41 were CD4^+^ T regulatory memory cells (T regs) and the P27 (unlabeled cells) were CD11c^+^ NK cells^36^. The up regulation of PD-L1 and CD86 in C2 was observed mostly in the immature granulocyte population of G5, G6 and G1, G5, G6, respectively and the low level of CD86 was also found in G4, CD11c^mid^ population (Figure 3A and C, left panel).

In general, the different diagnoses were distributed across clusters (Figure 3B) and CTR individuals were located within cluster C3, and less in C2, while no CTRs mapped into C1. These results show that patients can be classified based on the circulating cell phenotype, regardless of their diagnosis. The levels of surface proteins expressed by the cells give another important layer of information to stratify patients.

### 7.3 Mechanisms underlying patient reclassification

We next characterized the composition of the clusters using the clinical information (Figure 4A). Interestingly, we noticed a representation of each diagnostic group in all in 3 clusters. Treatments were also evenly distributed. Some enrichment for antimalarials in C1 cluster was observed compared to clusters C2 and C3 suggesting a more active phenotype in C1, although this was not significant. Accordingly, C4 hypocomplementemia was more frequent in cluster C1 compared to clusters C2 and C3, again pointing towards more active phenotype in C1. No differences in age or disease duration were found. Furthermore, no treatment influence on the cell frequency and MSI was observed besides that of CD25 expression in P27 being higher in the steroid-treated group (Table S3).

**Figure 4.**
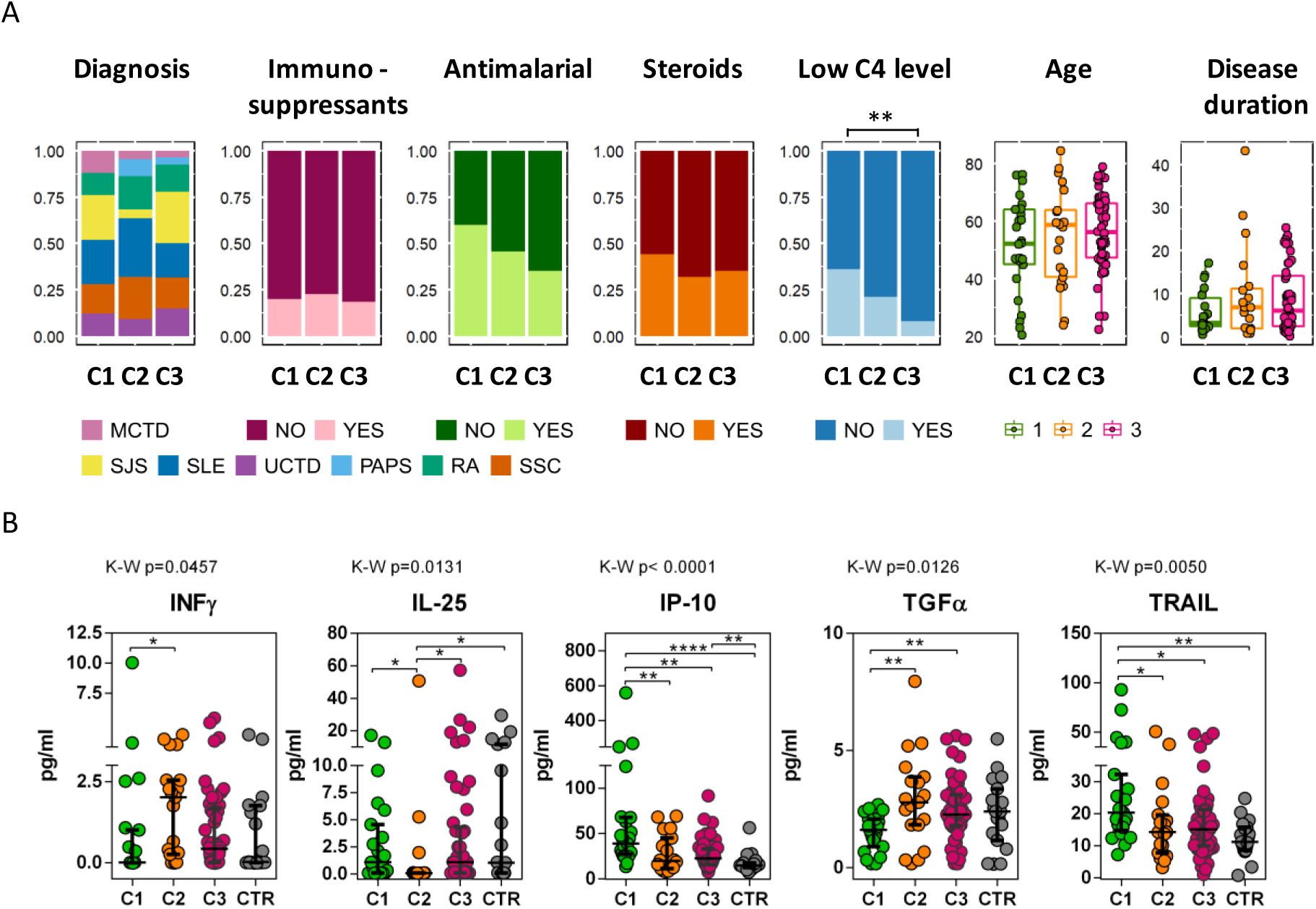
Patients in C1 present an inflammatory phenotype. (A) Distribution of different clinical parameters in the clusters identified in Figure 3 are represented on the x-axis. The y-axis in column plots represents the total frequency of individuals expressed as 1. Statistical analysis is performed using absolute numbers. Boxplots represents age and disease duration. Colors are represented as in the legend below. If parameters are statistically different between the groups p-value is drawn above the plot, Fisher exact test was used. (B) Cytokines level across identified clusters. Data are represented by the dot plot where the x-axis shows cluster labels together with CTRs group and the y-axis represents cytokine expression in pg/ml. Each dot represents one individual and is colored by the cluster colors, as in legend above. Differential analysis is performed using K-W followed by the Dunn test. The K-W p-values can be found above each plot and for the Dunn test statistical intervals are shown on the plots. The p-values for the Dunn test were adjusted using the FDR method to account for multiple comparisons. CTRs were not included in the statistical tests and are shown for reference purposes. Only cytokines with p-values ≤ 0.05 for K-W are shown. *p ≤ 0.05, **p ≤ 0.01, ***p ≤ 0.001 in Dunn test.

Since a common scale for disease activity across 7 different SADs does not exist, we measured the levels of 45 plasma cytokine many of which are known to be associated with disease activity (Figure 4B). The levels of IP-10 and TRAIL were elevated in cluster C1 as compared to C2 and C3, being higher also when compared to CTRs. These proteins were found to be associated with disease activity^37–39^ and interestingly induced by interferons^40,41^. It is worth mentioning that the level of TRAIL was also associated with antimalarial treatment (Table S4). Additionally, the level of the immunomodulatory cytokine TGFα was downregulated in cluster C1 as compared to C2, C3, and CTRs (Figure 4B). This supports the observation that C1 contains the least physiological and most active phenotype of patients.

On the other hand, the level of IL-25 (IL-17E), a cytokine that regulates inflammation by controlling the Th17 response, was down regulated in cluster C2 as compared to C1, C3, and CTRs. Both clusters C1 and C3 presented similar levels as CTRs (Figure 4B). Furthermore, patients in C2 showed the highest levels of IFNγ, especially when compared to C1. The down regulation of IL-25 upon regulation by IFNƴ may suggest a Th1/Th17 phenotype.

These results confirm the opposite composition and cellular function of cluster C1 and C2, suggesting different pathological mechanisms in these patients.

## 8 Discussion

In this study, we used in-depth immunophenotyping using mass cytometry and multiplex cytokine analysis to redefine groups of SADs patients. These groups seem to be related to different immunopathological mechanisms.

Firstly, using the classical approach, we compared different diagnostic entities and by clustering them we observed some similarities: SLE, SJS, and MCTD in group A1, RA, SSC, and UCTD in group A2, and CTR and PAPS in group A3. The close resemblance between SLE and SJS was already described using conventional cytometry^8^, however little is known about MCTD. MCTD is characterized by symptoms overlapping with those of SLE, but also of RA and SSc^42^. Interestingly, in terms of circulating cells phenotype, patients with MCTD were more similar to A1 (SLE, SJS) than to A2 (RA, SSC, UCTD). A similar behavior was also described using gene methylation analysis^43^, supporting our results, and emphasizing the notion that these diseases share immune pathological mechanisms. Remarkably, the most dysregulated feature present in group A1 was the elevated expression of CD38 in various cell types from the PBMC compartment: NK cells, naïve B cells, and DC. Of note, this multi-population dysregulation of CD38 was already reported in SLE^31^, and successful treatment of refractory SLE patients with Daratumumab (an anti-CD38) was described^32^. Hence, our data suggests that SJS and MCTD groups could potentially benefit from any line of treatment targeting CD38. SSC and UCTD patients have been reported to share clinical manifestations, and for this reason, the term UCTD-risk-SSC was developed to indicate that a high percentage of UCTD patients could develop SSC^44^. In a small cohort of patients, it was also observed that 90% of patients with SSC develop arthritis^45^, and consequently it may not be surprising that RA and SSC patients are included within group A2. Thus, our results suggest that UCTD patients could be closely monitored for risk of developing SSC or RA-related manifestations using whole blood immunophenotyping. Interestingly, controls separated from the patients, as expected, but fell into group A3 together with PAPS. Although clear differences could be observed between PAPS and CTRs, limited conclusions can be drawn due to the low number of PAPS patients.

In the second step of this study, patient clustering and reclassification analysis revealed 3 clusters of patients (C1 to C3). Clusters C1 and C2 seemed to have the most extreme phenotypes and C3 was less differentiated, representing, from our point of view, a transitional group between C1 and C2. We observed that features distinguishing overall SADs from CTRs were mostly dysregulated CD11c^+^ NK cells, CD4^+^ T reg cells, basophils, and the granulocyte compartment.

Although granulocyte/neutrophils are the predominant cells in the circulation, they are often discarded from the analyses. Neutrophils were historically considered as terminally differentiated, short-lived cells. However, the emerging evidence shows that neutrophils can survive much longer than previously thought, particularly under inflammatory conditions or through interaction with other cell subsets^46^. Neutrophils are well known for their involvement in SADs pathogenesis^47^ and the data pointed to a dysregulated expression of the cell death receptor Fas (CD95) in patients belonging to cluster C1. This may suggest neutrophil susceptibility to cell death, a process that is involved in autoimmunity^48^. In cluster C2 the level of CD95 was lower and similar to CTRs; however, high expression of the co-stimulatory molecules, PD-L1 and CD86 was observed. The expression of PD-L1 on neutrophils was already reported in SLE patients and associated with disease activity^49^. Our data show overexpression of PD-L1 in other diseases, extending the pathogenic role of this population to other SADs patients. Furthermore, reports have shown the ability of neutrophils to present antigens through MHC I/CD86 expression and to activate the adaptive immune response for example during infection^50^. Our results suggest that this process may have important implications in SADs and could be investigated further.

NK is a specialized compartment of innate lymphoid cells that has an essential role in initiating and maintaining the inflammatory and autoimmune response^51^. Here we report that patients belonging to cluster C1 had elevated expression of CD38 in CD56^hi^ CD11c^hi^ cells. This NK population was already described in SLE^31,52^, children with high risk for type I diabetes^53^ and in multiple sclerosis patients with clinical relapse^36^. Although no functional studies have been performed in humans, the evidence from mouse models suggests that these cells possess both NK-like and DC-like functions, being able to produce type I and II IFNs, perform antigen presentation, and trigger a lupus-like disease^51^. Our data highlight the need for further characterization of these NK cells in the context of the other SADs.

Another feature contributing to cluster C1 was CD25 expression in basophils. Basophils are known for their involvement in some allergic and parasitic diseases, however, their contribution to SLE and recently to MCTD was also reported^54^. CD25 (IL-2R) is expressed on effector T cells including T regs, however its role is not well explored in the context of basophils. A recent study showed that stimulation of human basophils with IL-2 triggers the expression of various proinflammatory cytokines^55^, suggesting that IL-2-sensitive basophils could participate in inflammatory responses. Since several clinical trials in various of the diseases studied here target IL-2/IL-2R, our data suggest that monitoring not only the T cell-mediated response but also basophils, could have important consequences in drug response or on therapeutic efficacy.

To gain more insight into the immunological status of the patients at the moment of sample collection, we analyzed cytokine production across the clusters. In cluster C1 we observed high levels of IP-10 and TRAIL and low levels of TGFα. IP-10 and TRAIL are upregulated by type I IFN^41^ and their level is elevated in some SAD patients, being also associated with disease activity in the case of IP-10, in SLE and RA patients^37,38^. Also, both TRAIL and CD95 belong to the TNFR superfamily and are involved in apoptosis, a process well-known to be implicated in SADs pathogenesis. This should indicate that the immunophenotype of C1 patients is associated with strong apoptosis and elevated IFN-I-related responses. These results are in agreement with their C4 hypocomplementemia and higher CD38 levels in CD25^+^ T cells, both associated with disease activity^56^. Patients in cluster C2 showed high IFNƴ levels together with low IL-25. IL-25 is an antagonist for IL-17A^57^, suggesting a Th1/Th17 driven phenotype. Interestingly, it has been shown that IFNƴ-treated neutrophils upregulate PD-L1 expression and acquire an immunosuppressive phenotype by limiting T cell proliferation^58^. Thus, it is possible that in this cluster the elevated PD-L1 expression is due to an enhanced Th1/Th17-mediated response. Since we did not observe any strong phenotype in cluster C3, we believe that this cluster represents less active patients that could progress either into C1 or C2-like phenotypes as represented in the UMAP visualization (Figure 3). A similar undefined group of patients was also reported by Barturen et al. This group gathered healthy controls and lower disease activity patients^9^.

Overall, we showed that by using mass cytometry and cytokine expression we can analyze patients suffering from different SADs at multiple cellular levels to define groups of patients sharing similar immune landscapes, who could benefit from the same line of treatment (see graphical abstract) Mass cytometry is not commonly used in the clinic, hence the translation of our results to the patients’ treatment and diagnosis is not straightforward. However, our data could serve as a basis for a simplified panel to be used in conventional cytometry, which could be translated to clinical practice and guide treatment decisions in a strategy closer to precision rheumatology.

## Acknowledgments

The research leading to these results has received support from the Innovative Medicines Initiative Joint Undertaking under grant agreement n° 115565 (PRECISESADS), resources of which are composed of financial contributions from the European Union’s Seventh Framework Programme (FP7/2007-2013) and EFPIA companies’ in-kind contribution, and funding from the Innovative Medicines Initiative 2 Joint Undertaking (JU) under grant agreement No 831434 (3TR). The JU receives support from the European Union’s Horizon 2020 research and innovation program and EFPIA. S.V.G. is an ISAC Marylou Ingram Scholar and is supported by an FWO postdoctoral research grant (1272823N, Research Foundation – Flanders). We are grateful to Olivia Santiago and Jose Diaz Cuéllar for technical support as a Core facility in Genyo. We would like to express our gratitude to the donors, your trust and health problems were the biggest motivation for this study. Special thanks to the very active community on cytoforum.stanford.edu, the mine of CyTOF-related information. We also acknowledge BioRender.com.

## 9 Data availability

Data will be available upon publication.

