## Supplementary material for "Systemic autoimmune disease patients’ blood immunome reveals specificities and commonalities among different diagnostic entities"

**Supplementary Figures**

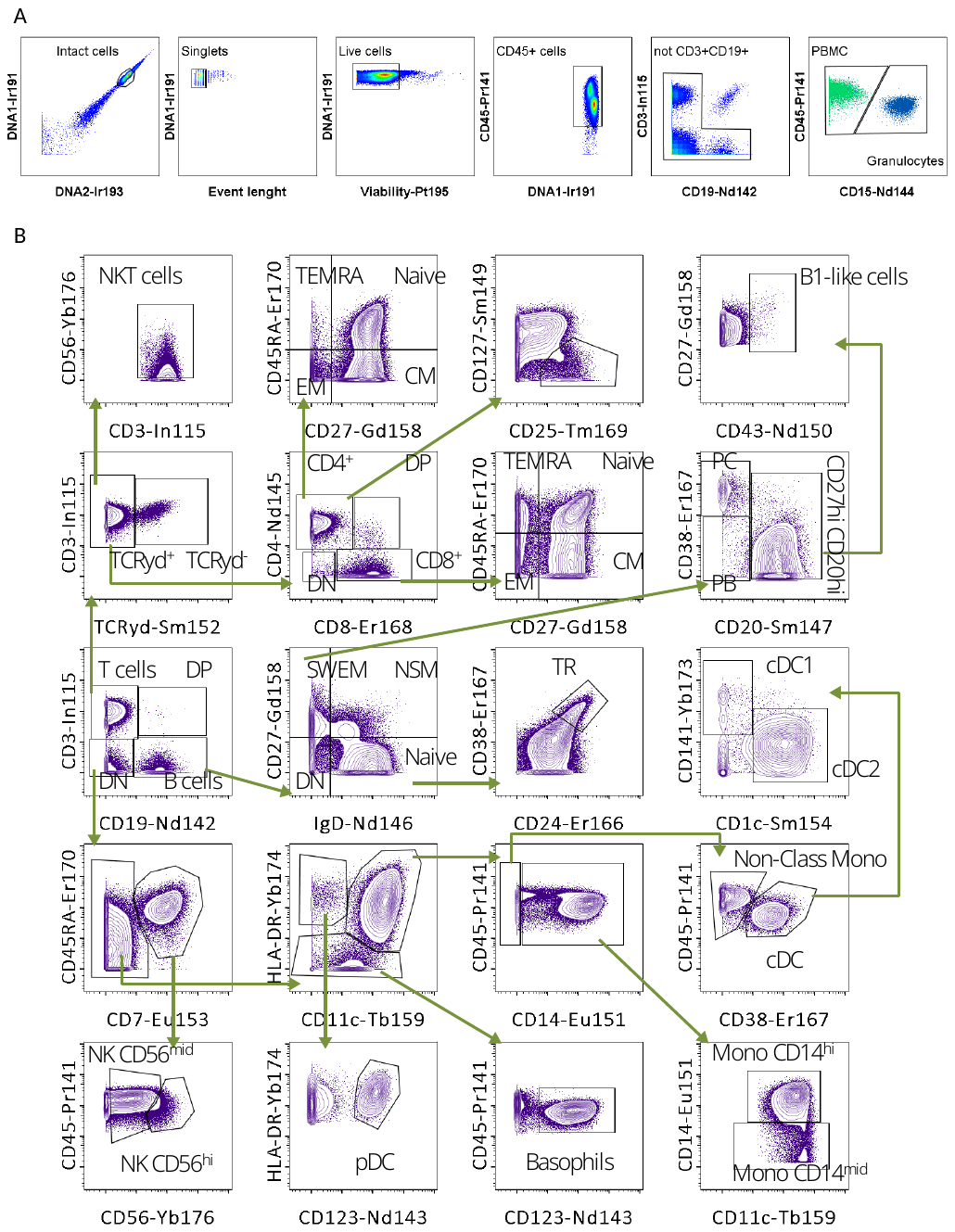

Figure S1 Gating. (A) Representative example gating strategy to obtain CD45^+^ cells and separating PBMC from GRAB using ratio computed for CD45, and CD15 markers. (B)The aggregated file was manually gated to obtain cluster and metacluster labels for FlowSOM tree. SWME – switched memory B cells, PC – Plasma cells, PB – Plasmablast, DP – double positive, DN – double negative, TR – Transitional B cells, TEMRA – Terminally differentiated T cells, CM – Central memory, EM – Effector memory, Mono – monocytes. Related to Figure 1.

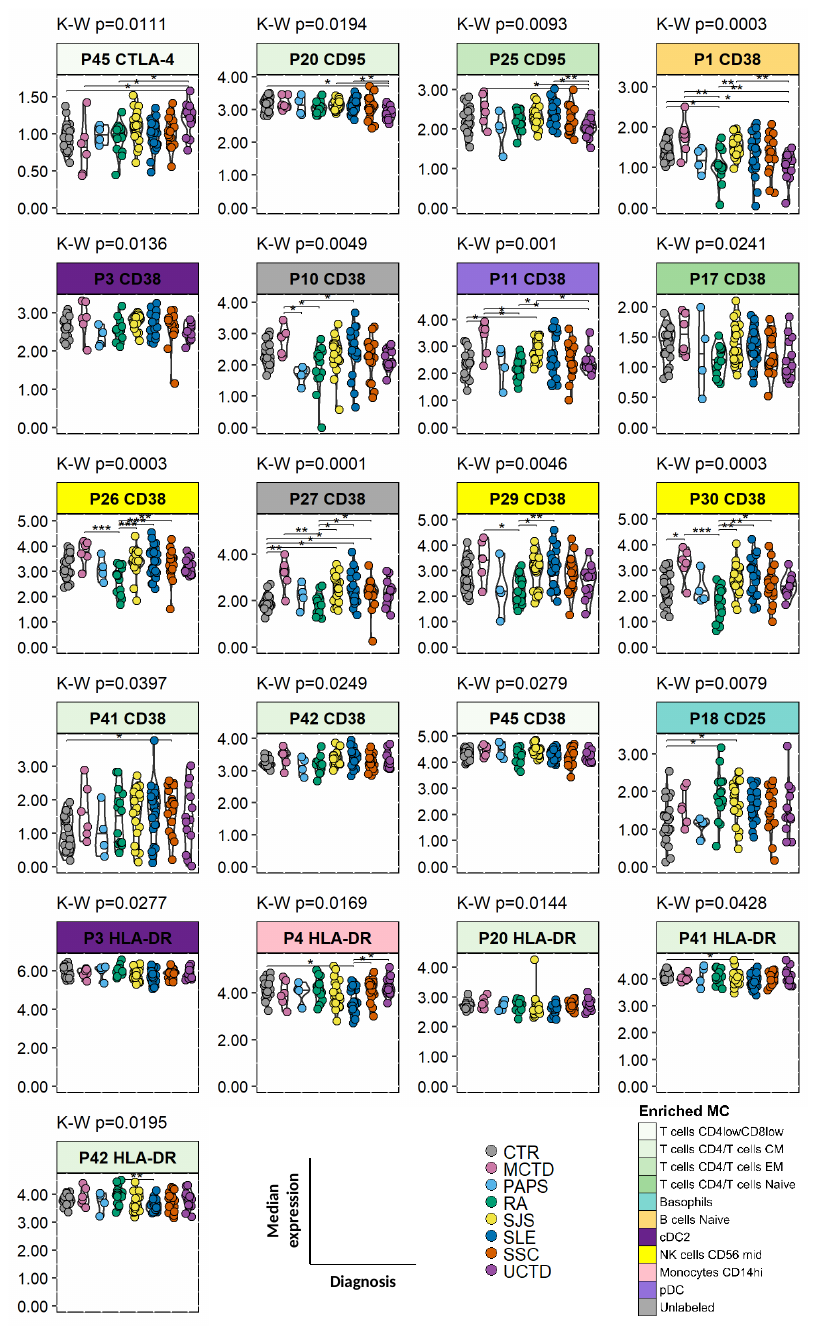

Figure S2 Multiple MSI in PBMC are differentially expressed between SADs. Cell frequencies were quantified for each metacluster as described in methods. Differential analysis was performed using K-W followed by a Dunn test. The K-W p-values can be found above each plot and the Dunn test statistical intervals are shown on the plots. p-values for the Dunn test were adjusted using the FDR method to account for multiple comparisons. Only metaclusters with p-value ≤ 0.05 for K-W are shown. Each plot is colored by the manual label of the metacluster and each dot in the plot represents an individual, colored by diagnosis. *p < 0.05, **p < 0.01, ***p < 0.001 in Dunn test. Related to Figure 2.

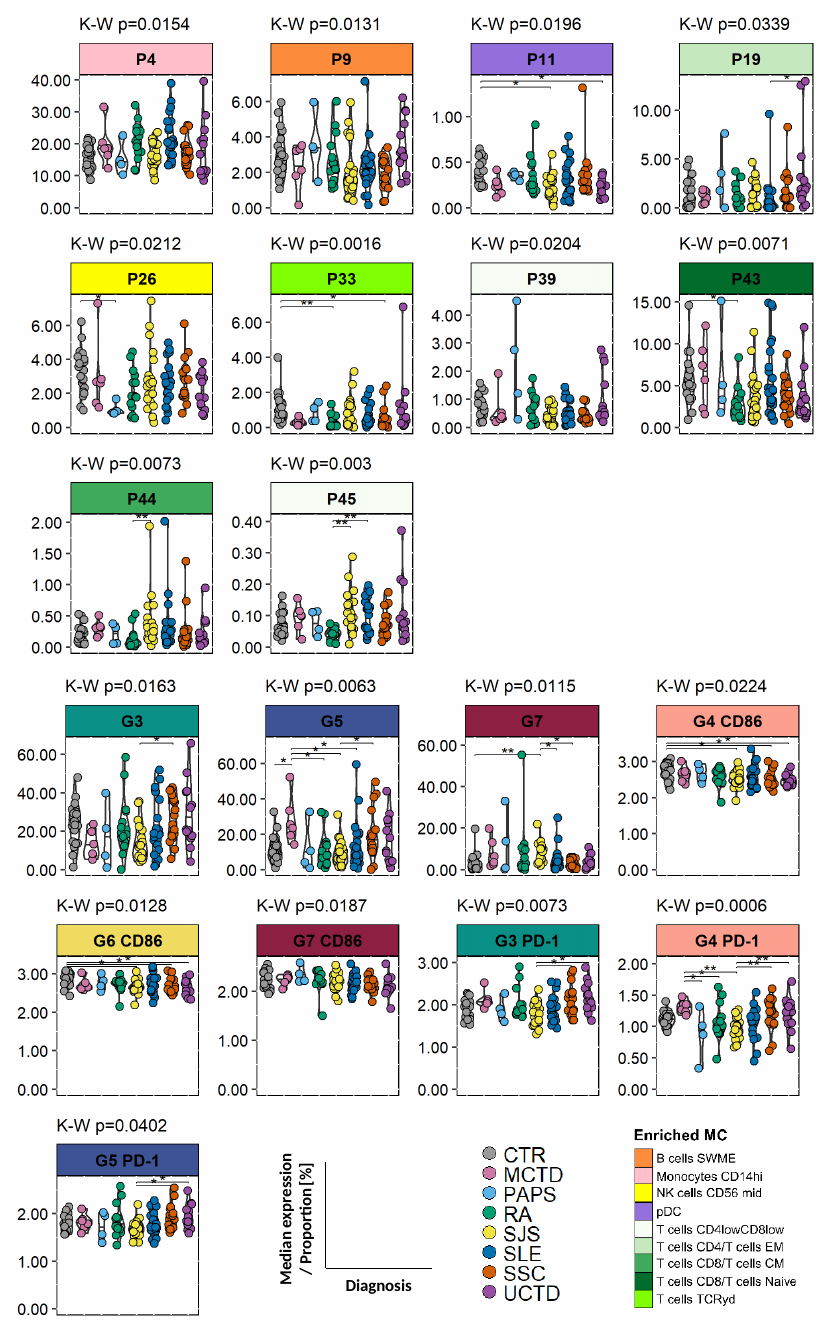

Figure S3 Various cell frequencies in PBMC and MSI with cell frequency in granulocytes are differentially expressed between SADs. Data are represented as in Figure S2. Related to Figure 2.

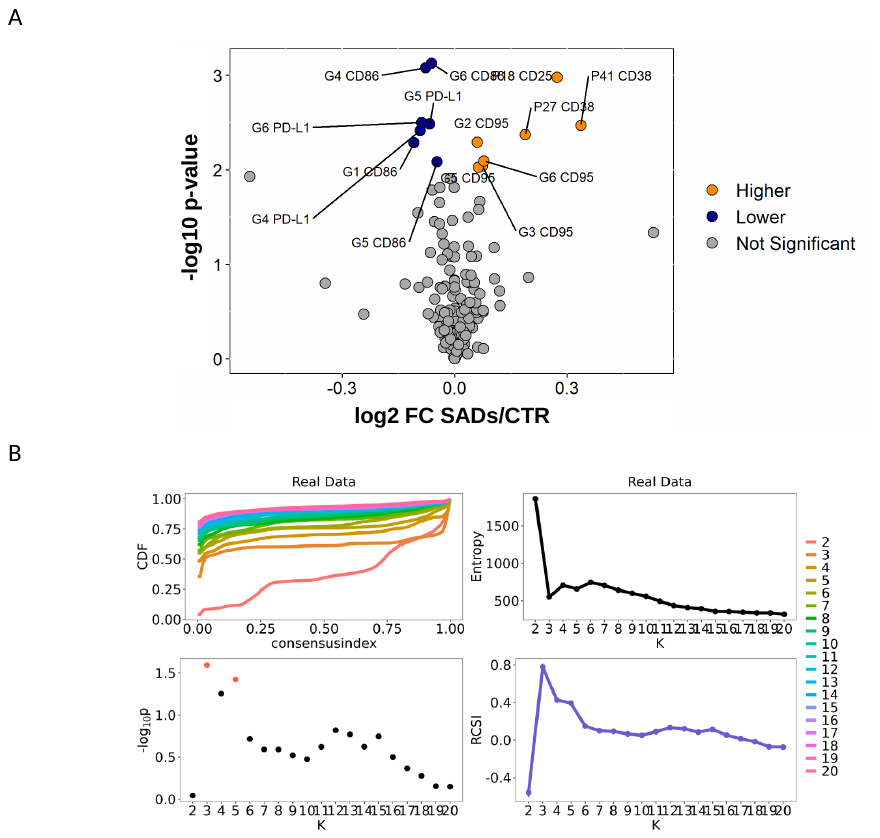

Figure S4 differentially express features separate patients in 3 stable clusters. (A) Volcano plot represents features being differentially expressed between SADs (autoimmune disease, N = 101) and CTR (control individuals, N = 22). The linear model was used. On the y-axis are the –log10 transformed p-values, and on the x-axis the log2 fold change (FC) between SADs and CTR. The blue and orange points represent features with lower or higher values in SADs than in CTR respectively, which are statistically significant (p-value ≤ 0.05). The grey points represent features that are not-differentially expressed. Features labeled only with P or G number represent frequency features, and those with P or G number and marker represents MSI. (B) Cluster stability using M3C method represented by a cumulative distribution function (CDF), entropy, p values and relative cluster stability index (RCSI) across 20 clusters (K). In the CDF line colors represent different K and in the p value plot, red dots represent significance. The p-values are represented as in A. Related to Figure 3.

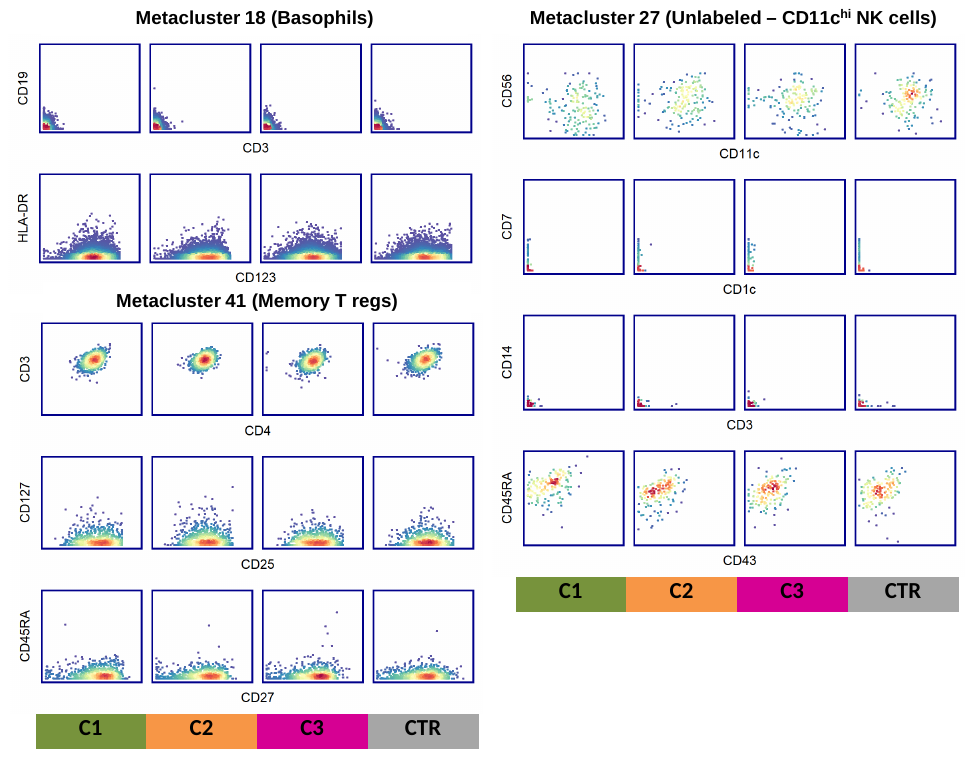

Figure S5 Autoimmune-specific metaclusters represent well-known cell populations. The aggregated file used to construct FlowSOM is used for visualization. Different markers are plotted to confirm metaclusters purity and its manual label assignment. Related to Figure 3.

**Supplementary Tables**

Table S1 Cohort demography and treatment

|  | CTR | MCTD | PAPS | RA | SJS | SLE | SSC | UCTD |
| --- | --- | --- | --- | --- | --- | --- | --- | --- |
| Age (years)^*^ | 46±7 | 44±12 | 40±17 | 64±10 | 57±10 | 45±15 | 60±14 | 60±16 |
| Female, n (%) | 22 (100) | 5 (83) | 2 (50) | 10 (67) | 20 (91) | 17 (74) | 17 (94) | 12 (92) |
| SAD duration (years)^*^ | NaN±NA | 8.7±6.9 | 9.7±8.5 | 6.4±6.1 | 8.3±8.4 | 8.7±7.2 | 11.2±11.6 | 4.3±4.5 |
| Antimalarials, n (%) | 0 (0) | 2 (33) | 1 (25) | 8 (53) | 8 (36) | 17 (74) | 2 (11) | 6 (46) |
| Immunosoppressants, n (%) | 0 (0) | 3 (50) | 0 (0) | 10 (67) | 3 (14) | 1 (4) | 3 (17) | 0 (0) |
| Steroids, n (%) | 0 (0) | 3 (50) | 0 (0) | 11 (73) | 6 (27) | 9 (39) | 4 (22) | 4 (31) |
| Antibiotics, n (%) | 0 (0) | 0 (0) | 0 (0) | 0 (0) | 0 (0) | 1 (4) | 2 (11) | 0 (0) |
| Center: COR, GRA, N | 0, 22 | 3, 3 | 0, 4 | 13, 2 | 8, 14 | 12, 11 | 13, 5 | 11, 2 |

CTRs: healthy controls, MCTD: mixed connective tissue disease, PAPS: primary antiphospholipid antibody syndrome, RA: rheumatoid arthritis, SJS: Sjögren’s syndrome, SLE: systemic lupus erythematosus, SSC: systemic sclerosis, UCTD: undifferentiated connective tissue disease, *mean ± SD in years, number of patients with assessable values, % = (n/N)*100.

Table S2 Antibody cocktail for high-content immunophenotyping

| Antigen | Clone | Metal | Source | Staining | Population definition |
| --- | --- | --- | --- | --- | --- |
| CD41 | HIP8 | 89Y | Standard BioTools | Clustering | Granulocytes |
| CD3 | UCHT1 | 115In | ThermoFisher | Clustering | PBMCs |
| CD45 | HI30 | 141Pr | Standard BioTools | Gating | - |
| CD19 | HIB19 | 142Nd | Standard BioTools | Clustering | PBMCs |
| CD123 | 6H6 | 143Nd | Standard BioTools | Clustering | PBMCs |
| CD15 | W6D3 | 144Nd | Standard BioTools | Clustering | Granulocytes |
| CD4 | RPA-T4 | 145Nd | Standard BioTools | Clustering | PBMCs |
| IgD | IA6-2 | 146Nd | Standard BioTools | Clustering | PBMCs |
| CD20 | 2H7 | 147Sm | Standard BioTools | Clustering | PBMCs |
| PD-L1 | 29E.2A3 | 148Nd | Standard BioTools | Functional | Granulocytes |
| CD127 | A019D5 | 149Sm | Standard BioTools | Clustering | PBMCs |
| CD43 | 84-3c1 | 150Nd | Standard BioTools | Clustering | PBMCs |
| CD14 | M5E2 | 151Eu | Standard BioTools | Clustering | PBMCs |
| TCRγδ | 11F2 | 152Sm | Standard BioTools | Clustering | PBMCs |
| CD7 | CD7-6B7 | 153Eu | Standard BioTools | Clustering | PBMCs |
| CD1c | L161 | 154Sm | Biolegend | Clustering | PBMCs |
| BAFF-R | 11C1 | 155Gd | Standard BioTools | Functional | - |
| CD86 | IT2.2 | 156Gd | Standard BioTools | Clustering / Functional | Granulocytes |
| CD27 | L128 | 158Gd | Standard BioTools | Clustering | PBMCs |
| CD11c | Bu15 | 159Tb | Standard BioTools | Clustering | PBMCs |
| CD28 | CD28.2 | 160Gd | Standard BioTools | - | - |
| CTLA-4 | 14D3 | 161Dy | Standard BioTools | Functional | - |
| CD69 | FN50 | 162Dy | Standard BioTools | Functional | - |
| CD95 | DX2 | 164Dy | Standard BioTools | Clustering / Functional | Granulocytes |
| CD40 | 5C3 | 165Ho | Standard BioTools | Functional | - |
| CD24 | ML5 | 166Er | Standard BioTools | Clustering | Both |
| CD38 | HIT2 | 167Er | Standard BioTools | Clustering / Functional | PBMCs |
| CD8 | SK1 | 168Er | Standard BioTools | Clustering | PBMCs |
| CD25 | 2A3 | 169Tm | Standard BioTools | Clustering / Functional | PBMCs |
| CD45RA | HI100 | 170Gd | Standard BioTools | Clustering | PBMCs |
| CD57 | HCD57 | 172Yb | Standard BioTools | Clustering | PBMCs |
| CD141 | 1A4 | 173Yb | Standard BioTools | Clustering | PBMCs |
| HLA-DR | L243 | 174Yb | Standard BioTools | Clustering /Functional | PBMCs |
| PD-1 | EH12.2H7 | 175Lu | Standard BioTools | Clustering / Functional | Both |
| CD56 | N901 | 176Yb | Standard BioTools | Clustering | PBMCs |
| CD16 | 3G8 | 209Bi | Standard BioTools | - | - |
| CisPt | Live/dead cells | 195Pt | Sigma | Intracellular | - |
| DNA1 | Nucleated cells | 191Ir | Standard BioTools | Intracellular | - |
| DNA2 | Nucleated cells | 193Ir | Standard BioTools | Intracellular | - |
| Bead | Beads | 140Ce | Standard BioTools | - | - |

Table S3 Treatment influence on cell frequency and MSI

| **Median [IQR]** | | | | **Mann-Whitney [FDR]** | | |
| --- | --- | --- | --- | --- | --- | --- |
| **Populations** | **CTRs** | **Tx** | **NTx** | **CTRs-NTx** | **CTRs-Tx** | **NT-Tx** |
| **Antimalarials CTRs (N = 22), Tx (N = 44), NTx (N = 57)** | | | | | | |
| G1_CD86 | 2.14 [2.01 – 2.23] | 1.87 [1.7 – 2.01] | 1.89 [1.73 – 2.07] | ** | ** | ns |
| G2_CD95 | 1.97 [1.93 – 2.08] | 2.13 [2.04 – 2.32] | 2.09 [1.99 – 2.24] | * | ** | ns |
| G3_CD95 | 1.93 [1.77 – 2.02] | 2.05 [1.96 – 2.29] | 2.01 [1.89 – 2.21] | ns | ** | ns |
| G4_CD86 | 2.79 [2.64 – 2.91] | 2.49 [2.39 – 2.73] | 2.57 [2.39 – 2.75] | ** | ** | ns |
| G4_PD_L1 | 1.98 [1.83 – 2.09] | 1.76 [1.56 – 1.88] | 1.75 [1.63 – 1.9] | ** | ** | ns |
| G5_CD86 | 3.17 [2.97 – 3.26] | 2.93 [2.82 – 3.12] | 2.96 [2.84 – 3.08] | * | * | ns |
| G5_CD95 | 1.94 [1.81 – 2.01] | 2.06 [1.98 – 2.27] | 2.05 [1.87 – 2.16] | ns | ** | ns |
| G5_PD_L1 | 2.3 [2.19 – 2.44] | 2.07 [1.97 – 2.26] | 2.13 [2.05 – 2.29] | * | ** | ns |
| G6_CD86 | 2.89 [2.74 – 2.98] | 2.68 [2.58 – 2.83] | 2.69 [2.58 – 2.81] | *** | ** | ns |
| G6_CD95 | 1.67 [1.57 – 1.77] | 1.86 [1.73 – 2.08] | 1.83 [1.64 – 1.97] | * | ** | ns |
| G6_PD_L1 | 2.04 [1.9 – 2.13] | 1.85 [1.73 – 2.05] | 1.83 [1.66 – 2] | ** | * | ns |
| P18_CD25 | 1.15 [0.92 – 1.47] | 1.72 [1.43 – 2.08] | 1.5 [1.15 – 2] | * | ** | ns |
| P27_CD38 | 1.85 [1.73 – 2.13] | 2.36 [1.91 – 2.98] | 2.34 [2.01 – 2.58] | ** | ** | ns |
| P41_CD38 | 0.89 [0.55 – 1.42] | 1.84 [1.2 – 2.12] | 1.6 [0.73 – 2.17] | * | *** | ns |
| **Immunosuppressants CTRs (N = 22), Tx (N = 20), NTx (N = 81)** | | | | | | |
| G1_CD86 | 2.14 [2.01 – 2.23] | 1.82 [1.7 – 2.08] | 1.89 [1.73 – 2.06] | ** | * | ns |
| G2_CD95 | 1.97 [1.93 – 2.08] | 2.11 [1.98 – 2.22] | 2.11 [2.04 – 2.28] | ** | ns | ns |
| G3_CD95 | 1.93 [1.77 – 2.02] | 2.02 [1.88 – 2.22] | 2.03 [1.93 – 2.24] | * | ns | ns |
| G4_CD86 | 2.79 [2.64 – 2.91] | 2.53 [2.41 – 2.75] | 2.51 [2.39 – 2.73] | ** | * | ns |
| G4_PD_L1 | 1.98 [1.83 – 2.09] | 1.71 [1.59 – 1.91] | 1.76 [1.61 – 1.88] | ** | * | ns |
| G5_CD86 | 3.17 [2.97 – 3.26] | 2.97 [2.89 – 3.08] | 2.96 [2.8 – 3.1] | * | ns | ns |
| G5_CD95 | 1.94 [1.81 – 2.01] | 2.08 [1.88 – 2.19] | 2.06 [1.91 – 2.22] | * | ns | ns |
| G5_PD_L1 | 2.3 [2.19 – 2.44] | 2.12 [1.99 – 2.31] | 2.11 [2.02 – 2.26] | ** | ns | ns |
| G6_CD86 | 2.89 [2.74 – 2.98] | 2.72 [2.61 – 2.81] | 2.68 [2.56 – 2.82] | *** | * | ns |
| G6_CD95 | 1.67 [1.57 – 1.77] | 1.84 [1.54 – 2.02] | 1.83 [1.7 – 2.01] | ** | ns | ns |
| G6_PD_L1 | 2.04 [1.9 – 2.13] | 1.85 [1.67 – 2.03] | 1.84 [1.69 – 1.99] | ** | ns | ns |
| P18_CD25 | 1.15 [0.92 – 1.47] | 1.7 [1.18 – 1.98] | 1.56 [1.27 – 2.01] | ** | ns | ns |
| P27_CD38 | 1.85 [1.73 – 2.13] | 2.1 [1.61 – 2.73] | 2.37 [2.01 – 2.79] | *** | ns | ns |
| P41_CD38 | 0.89 [0.55 – 1.42] | 1.61 [0.73 – 2.1] | 1.76 [1.01 – 2.17] | ** | * | ns |
| **Steroids CTRs (N = 22), Tx (N = 37), NTx (N = 64)** | | | | | | |
| G1_CD86 | 2.14 [2.01 – 2.23] | 1.89 [1.74 – 2.03] | 1.87 [1.69 – 2.07] | ** | ** | ns |
| G2_CD95 | 1.97 [1.93 – 2.08] | 2.15 [2.04 – 2.32] | 2.09 [2.02 – 2.22] | ** | ** | ns |
| G3_CD95 | 1.93 [1.77 – 2.02] | 2.06 [1.91 – 2.35] | 2.01 [1.93 – 2.16] | * | * | ns |
| G4_CD86 | 2.79 [2.64 – 2.91] | 2.47 [2.38 – 2.73] | 2.53 [2.4 – 2.73] | ** | ** | ns |
| G4_PD_L1 | 1.98 [1.83 – 2.09] | 1.75 [1.59 – 1.93] | 1.77 [1.63 – 1.9] | ** | ** | ns |
| G5_CD86 | 3.17 [2.97 – 3.26] | 2.97 [2.81 – 3.12] | 2.95 [2.84 – 3.07] | * | * | ns |
| G5_CD95 | 1.94 [1.81 – 2.01] | 2.09 [1.87 – 2.26] | 2.06 [1.94 – 2.18] | * | * | ns |
| G5_PD_L1 | 2.3 [2.19 – 2.44] | 2.1 [1.99 – 2.27] | 2.14 [2.02 – 2.29] | ** | ** | ns |
| G6_CD86 | 2.89 [2.74 – 2.98] | 2.68 [2.58 – 2.82] | 2.69 [2.58 – 2.81] | ** | ** | ns |
| G6_CD95 | 1.67 [1.57 – 1.77] | 1.91 [1.7 – 2.1] | 1.8 [1.66 – 1.94] | * | * | ns |
| G6_PD_L1 | 2.04 [1.9 – 2.13] | 1.8 [1.64 – 1.98] | 1.87 [1.71 – 2] | * | ** | ns |
| P18_CD25 | 1.15 [0.92 – 1.47] | 1.77 [1.37 – 2.22] | 1.52 [1.18 – 1.97] | * | ** | * |
| P27_CD38 | 1.85 [1.73 – 2.13] | 2.23 [1.97 – 2.81] | 2.41 [1.98 – 2.71] | ** | ** | ns |
| P41_CD38 | 0.89 [0.55 – 1.42] | 1.66 [0.75 – 2.09] | 1.78 [1 – 2.18] | ** | * | ns |

The median and interquartile ranges are shown for each marker expression across CTR – healthy control, Tx – treated, NTx – non-treated. MW analysis is performed for each medication between the groups as indicated in the table. P values are interpreted as follows ns p > 0.05, *p < 0.05, **p < 0.01, ***p < 0.001

Table S4 Treatment influence on cytokine production

| **Median [IQR]** | | | | **Mann-Whitney [FDR]** | | |
| --- | --- | --- | --- | --- | --- | --- |
| Cytokines | CTRs | **Tx** | **NTx** | **CTR-NTx** | **CTR-Tx** | **NTx-Tx** |
| **Antimalarial CTR (N = 19), Tx (N = 41), NTx (N = 51)** | | | | | | |
| IFNγ | 0.03 [0.01 - 1.65] | 0.27 [0 - 1.08] | 0.63 [0.01 - 2.09] | ns | ns | ns |
| IL-25 | 1.04 [0.09 - 10.64] | 0.33 [0.09 - 5.88] | 0.09 [0.08 - 1.69] | ns | ns | ns |
| IP-10 | 14.71 [12.47 - 17.08] | 24.02 [15.45 - 38.9] | 26.82 [17.96 - 44.33] | *** | ** | ns |
| TGFα | 2.4 [1.32 - 3.12] | 2.07 [1.3 - 3.08] | 2.27 [1.68 - 2.91] | ns | ns | ns |
| TRAIL | 11.19 [9.13 - 15.52] | 18.56 [14.25 - 25.45] | 14.26 [9.78 - 19.73] | ns | ** | * |
| **Immunosuppressants CTRs (N = 19), Tx (N = 19), NTx (N = 73)** | | | | | | |
| IFNγ | 0.03 [0.01 - 1.65] | 0.01 [0.01 - 0.84] | 0.46 [0.01 - 2.02] | ns | ns | ns |
| IL-25 | 1.04 [0.09 - 10.64] | 0.09 [0.08 - 0.66] | 0.55 [0.08 - 4.99] | ns | ns | ns |
| IP-10 | 14.71 [12.47 - 17.08] | 22.74 [17.62 - 50.01] | 26.06 [16.09 - 43.62] | *** | *** | ns |
| TGFα | 2.4 [1.32 - 3.12] | 2.27 [1.62 - 2.78] | 2.07 [1.3 - 2.93] | ns | ns | ns |
| TRAIL | 11.19 [9.13 - 15.52] | 17.23 [12.74 - 24.12] | 16.02 [10.3 - 21.73] | * | * | ns |
| **Steroids CTRs (N = 19), Tx (N = 36), NTx (N = 56)** | | | | | | |
| IFNγ | 0.03 [0.01 - 1.65] | 0.28 [0.01 - 1.1] | 0.45 [0.01 - 2.06] | ns | ns | ns |
| IL-25 | 1.04 [0.09 - 10.64] | 0.11 [0.08 - 3.52] | 0.17 [0.08 - 3.46] | ns | ns | ns |
| IP-10 | 14.71 [12.47 - 17.08] | 30.09 [19.35 - 49.85] | 23.01 [15.46 - 34.57] | ** | *** | ns |
| TGFα | 2.4 [1.32 - 3.12] | 2.37 [1.48 - 3.52] | 2.06 [1.39 - 2.64] | ns | ns | ns |
| TRAIL | 11.19 [9.13 - 15.52] | 17.45 [12.95 - 22] | 15.33 [10.35 - 21.88] | ns | * | ns |

The median and interquartile ranges are shown for each marker expression across CTR – healthy control, Tx – treated, NTx – non-treated. MW analysis is performed for each medication between the groups as indicated in the table. P values are interpreted as follows ns p > 0.05, *p < 0.05, **p < 0.01, ***p < 0.001
